# Robustness of local control strategies under modelling uncertainties: the 2004 avian influenza H5N1 outbreak in Thailand

**DOI:** 10.1101/355024

**Authors:** Renata Retkute, Chris P. Jewell, Thomas P. Van Boeckel, Geli Zhang, Xiangming Xiao, Weerapong Thanapongtharm, Matt Keeling, Marius Gilbert, Michael J. Tildesley

## Abstract

The Highly Pathogenic Avian Influenza (HPAI) subtype H5N1 virus persists in many countries and has been circulating in poultry, wild birds. In addition, the virus has emerged in other species and frequent zoonotic spillover events indicate that there remains a significant risk to human health. It is crucial to understand the dynamics of the disease in the poultry industry to develop a more comprehensive knowledge of the risks of transmission and to establish a better distribution of resources when implementing control. In this paper, we develop a set of mathematical models that simulate the spread of HPAI H5N1 in the poultry industry in Thailand, utilising data from the 2004 epidemic. The model that incorporates the intensity of duck farming when assessing transmision risk provides the best fit to the spatiotemporal characteristics of the observed outbreak, implying that intensive duck farming drives transmission of HPAI in Thailand. We also extend our models using a sequential model fitting approach to explore the ability of the models to be used in “real time” during novel disease outbreaks. We conclude that, whilst predictions of epidemic size are estimated poorly in the early stages of disease outbreaks, the model accurately predicts the preferred control policy that should be deployed to minimise the impact of the disease.

## 1. Introduction

Since the emergence of highly pathogenic avian influenza (HPAI) H5N1 in the late 1990s, the virus has had significant impact on poultry industries around the world and posed a serious threat to public health. The majority of outbreaks occur in East and South Asia, a region that contains half of the population of the world. Whilst the probability of a human AI pandemic may appear low based on the poor ability of the virus to adapt to the upper human airway (Peiris et al. 2007), the death toll of such an event may be catastrophic. As a result of the HPAI H5N1 panzootic, there have been 860 confirmed human cases of avian influenza A (H5N1) in 16 countries and 454 deaths (a human case fatality of 53%) as of 30th October, 2017, based on World Health Organization (WHO) statistics. The majority of humans infected with the virus work in professions that involve close contact with potentially infected poultry (de Bruin et al. 2017) and therefore it is crucial to understand the dynamics of the disease in the poultry industry to develop a more comprehensive knowledge of the risks of transmission to humans.

In Thailand, H5N1 was first detected in the poultry industry in January 2004 (Tiensin et al. 2005). The first wave of infection took place from January to May and resulted in 193 reported AI outbreaks. A second infection wave started in July 2004 and culminated in March 2005, with 1,492 outbreaks notified during this period. Approximately 62 million birds were killed either through infection or through targeted culling as a control measure. In addition, during the 2004 H5N1 epidemic there were 17 human cases of infection and 12 deaths (Tiensin et al. 2005, Auewarakul 2008). In an attempt to improve case detection, the Thai government implemented an X-ray survey (Gilbert et al. 2006) in September 2004. Control policies such as localised movement restrictions and 1km ring culling were introduced to reduce the risk of further spread of the disease; vaccination was not used.

The spatial distribution of HPAI H5N1 has been studied in several countries and regions using spatial statistical models (Gilbert et al. 2007, Truscott et al. 2007, Sharkey et al. 2008, Gilbert et al. 2008, Jewell et al. 2009, Gilbert et al. 2010, Stegeman et al. 2011, Ssematimba et al. 2012, Paul et al. 2014, Gilbert et al. 2014, Artois et al. 2017) and mathematical models (Minh et al. 2011, Walker et al. 2012, Hill et al. 2017) in East Asia. Evidence from previous work suggests that different landscapes, production systems and water-related variables are risk factors that will promote disease transmission. In addition, mathematical models such as the one that we present here can improve understanding about the transmission dynamics of infectious diseases and help with an assessment of the effectiveness of control strategies applied during outbreaks (Stegeman et al. 2011). A key challenge when implementing control is the lack of clinical signs in many duck species - whilst chickens have a very high mortality rate for H5N1 (Yu et al. 2007, Jeong et al. 2009), ducks are largely asymptomatic but can readily transmit the disease. In addition, in Thailand, as in many other countries in South East Asia, the presence of free grazing ducks, that feed year round on rice paddies, have been shown to be a strong predictor for the presence of the disease in the landscape (Gilbert et al. 2007, 2008). It is therefore crucial to develop a more detailed understanding of the factors leading to persistence of AI in poultry, the conditions that are most suitable to disease transmission, and intervention strategies that will minimize the future impact of the disease.

In this paper, we develop a set of mathematical models to simulate the spread of HPAI H5N1 in the poultry industry in Thailand. We use detailed demographic and epidemiological data from the 2004 epidemic and fit our models to epidemiological surveillance (X-Ray) data. Our framework will allow us to establish the risk factors that result in transmission of the disease that will in turn enable us to effectively target intervention strategies. We will also consider the ability of the models to be used in “real time”, whereby model parameters are obtained by only using data that are observable at a given stage of the epidemic. This enables us to determine the predictive capacity of HPAI models both in terms of establishing transmission risk and in determining the most appropriate intervention policy. It also allows us to quantify how progressive accumulation of information shapes the knowledge of outbreak dynamics and the robustness of control strategies.

## 2. Materials and methods

### 2.1. Data

In this paper, we use demographic data compiled by the Department of Livestock Development (DLD, Bangkok) that was constructed through the X-Ray surveys that were implemented in response to the H5N1 outbreak in 2004. Demographic data was recorded at the farm/owner level and consists of the number of chicken and ducks in each flock, and a unique identifier of the sub-district where each farm is located.

Owing to imperfect reporting at the onset of the epidemic, data for the first wave of infection in Thailand are largely incomplete. Therefore, for parameter estimation and analysis, we use the outbreak data from the second wave of the epidemic that took place from July 2004. In this second wave, more than 1400 outbreaks were reported and over 62 million birds died or were culled as part of the control policy (Tiensin et al. 2005). The outbreak data set includes the unique identifier of the subdistrict as well as the total number of sick, dead and destroyed animals.

In order to implement our mathematical model, we use geographical data for each subdistrict including the locations of subdistrict boundaries and the area of each subdistrict. We infer indices of neighbouring subdistricts by analysing intersection sets of subdistrict boundaries. We then calculate the length of all shared boundaries by fitting a linear spline through shared points on boundaries of neighbour subdistricts. The proportion of area used by rice paddy fields within each subdistrict was derived from the moderate resolution imaging spectroradiometer (MODIS) sensor onboard the NASA Terra satellite (Xiao et al. 2006).

### 2.2. Mathematical models

We develop a meta-population framework for our model, following the approach of Buhnerkempe et al. 2014. In our model, the basic unit of infection is the flock, and we assume that all birds (chicken and ducks) within a flock become infected such that an entire flock is classified as Susceptible, Exposed, Infectious or Removed.

In our modelling framework, we consider transmission via one of five mechanisms: (i) spatially independent transmission to any subdistrict in Thailand, (ii) transmission within an subdistrict, (iii) transmission across borders to neighbouring subdistricts, (iv) transmission driven by presence of rice paddies and (v) transmission driven by duck farm intensity. By varying the combination of these factors, we have formulated four mathematical models with increasing complexity: a random process model (Model A), a spatial model (Model B), a spatial model incorporating rice density (Model C), and a spatial model incorporating duck intensity (Model D). Details of the equations governing each model are given in the Supplementary Information.

#### 2.2.1. Model A: Random process model

The simplest model for a HPAI outbreak is that of complete spatial randomness, whereby transmission events are distributed independently according to a uniform probability distribution over all susceptible flocks. Under this scenario, the infection pressure at time *t* in any susceptible flock is equal to the background term, *δ*, multiplied by the number of infectious flocks at time *t*, *n*_*I*_(*t*), plus one. The biological meaning of the model is that the outbreak dynamics does not depend on any environmental factors or the structure of the poultry sector in Thailand. This naive model, whilst unlikely to be representative of heterogenous transmission routes, provides a baseline for epidemic spread for comparison with the spatially explicit models.

#### 2.2.2. Model B: Spatial model

The Spatial model utilises a classical metapopulation approach, based upon the model first used by Buhnerkempe et al. 2014. This model is an extension of the random process model, with the addition of within and between subdistrict transmission. The model therefore incorporates local density-dependent spread and contains three additive terms representing the different transmission scenarios - the background term, *δ*, within-subdistrict transmission with transmission rate *β*_*W*_ and local cross-border transmission with transmission rate *β*_*B*_. The framework assumes that transmission is dependent upon the poultry industry features within a given subdistrict and within neighbouring subdistricts and ignores the presence of other landscape features that may results in increased transmission risk. Within and between subdistrict transmission depends on the structure of poultry industry, and is parametrised by *p*_*C*_ and *p*_*D*_ (power laws for a number of chicken and ducks) and *ξ* (multiplier for relative infectiousness and transmisability of ducks to chicken).

#### 2.2.3. Model C: Spatial (rice) model

Given that previous work suggests that the presence of free grazing ducks can result in increased transmission risk in Thailand (Gilbert et al. 2007), here we use data on rice fields in the country (Xiao et al. 2006) and extend the meta-population model described in the spatial model above to consider the likelihood of increased transmission risk in subdistricts with a high density of rice paddies. In order to account for the role of rice paddies in the infection dynamics, we use a sigmoidal function that operates as a scaling factor for the infection pressure. Here a higher infection pressure is presumed to be present for subdistricts with a higher proportion of rice paddy fields. This dependence is described by the scaling, *α*_*R*_, threshold, *∊*_*R*_ and power, *n*_*R*_. Details of the equation are given in the Supplementary Information.

#### 2.2.4. Model D: Spatial (duck) model

Our final model builds upon the spatial model but additionally considers the intensity of duck farming within a subdistrict as an additional factor influencing transmission. We therefore categorise subdistricts based upon the intensity of their duck farming industry defined as a log_10_ of the size of largest duck flock within a subdistrict plus one. Furthermore, in a similar way as for the spatial (rice) model, we use a sigmoidal function such that a higher infection pressure is assumed for subdistricts with more intense duck farming. This dependence is described by the scaling, *α*_*D*_, threshold, *∊*_*D*_ and power, *n*_*D*_.

### 2.3. Fitting the models to outbreak data

Having defined the full modelling framework, we will consider this suite of nested mathematical models (A-D) to determine the factors that contribute to transmission of HPAI H5N1 within Thailand. Each model is fitted to outbreak data using a Bayesian likelihood approach that determines the parameters that best captures the observations (for details see Supplementary Material section below).

For a sequential analysis, we re-estimated model parameters by inclemently increasing the length of the outbreak data, evaluating how increasing amounts of outbreak data altered and influenced the parameter estimation and model projections.

### 2.4. Control options

Once fully parameterised, we will use the models to investigate the effectiveness of a range of interventions that could be implemented to control the outbreak. We will analyse outbreak projections under three competing strategies: (i) culling of identified infected premises (IP) only, (ii) culling of IPs and farms within a given radius of IPs (ring culling) and (iii) IP culling and reactive vaccination with a given capacity within infected subdistricts.

The spatially aggregate nature of the meta-population model that we have developed prevents us from applying a culling radius directly to each IP to determine which flocks will be culled. In order to mimic ring culling in our metapopulation framework we therefore reduce the number of susceptible flocks within an infected subdistrict as a fraction of the subdistrict area to the area of a circle with a particular culling radius. We analyse projections based on the following culling radius values: 0.25km, 0.5km, 1km, 2km and 5km.

As an alternative to ring culling, we explore the impact of reactive vaccination within any subdistrict reporting infection. For this control policy, following the removal of infected flocks, we assume that a given percentage of the poultry within an infected subdistrict are vaccinated and become immune to infection. We make the assumption that any poultry that are targeted for vaccination will acquire immunity 7 days after the relevant infected premises is reported. We then explore the impact of vaccination upon the outbreak size and duration, given vaccine capacities of 50%, 70% and 90% of all poultry within each infected subdistrict. See Supplementary Information for more details. As optimal control actions for control of disease outbreaks may depend on objectives (Probert et al. 2016), we evaluated the effectiveness of local control by analysing projected number of infected flocks, number of culled flocks and duration of outbreak.

## 3. Results

### 3.1. Model Parameterisation

We used a Bayesian framework to parameterise each model to the 2004 epidemics of H5N1 in Thailand. In order to compare the four modelling frameworks and establish a goodness of fit to the observational data, we calculate the deviance information criterion (DIC). Model with a lower DIC value implies a closer fit to the observational data compared to the alternative models and therefore this model is preferred. For DIC calculations we use the parameter values generated from our MCMC runs for parameter estimation (Spiegelhalter et al. 2002). We calculated the following values for the four models: *DIC*_*A*_ = 37478.2, *DIC*_*B*_ = 33763.5, *DIC*_*C*_ = 33596.7 and *DIC*_*D*_ = 33164.3.

The DIC values for each model indicate that Model D, incorporating the intensity of the duck industry, is preferred over all other modelling frameworks, indicating that it is the intensive duck farming industry that is predominantly driving transmission in Thailand.

For all models, the fitted posterior mean and 95% credible intervals for estimated parameters are summarised in Table 1. As the complexity of the modelling framework increases, the value of the background term *δ* decreases, as local scale characteristics drive transmission for the more complex models. When considering rice paddy density as a risk factor (Model C) similar parameter estimates are found for the background term and within-subdistrict transmission rate when compared with the purely spatial Model B. Interestingly Model D, that includes intensive duck farms as a risk factor, produces lower parameter value estimates for *δ*, *β*_*W*_ and *β*_*B*_, suggesting that the presence of intensive duck farms may have a strong influence on transmission.

**Table 1.** Estimated parameters with posterior mean and 95% CI for random process model (A), spatial model (B), spatial rice model (C), and spatial duck model (D).

| Parameter | Model (A) | Model (B) | Model (C) | Model (D) |
| --- | --- | --- | --- | --- |
| $\delta \times 10^{-8}$ | 2.55 (2.43, 2.68) | 1.23 (1.14, 1.30) | 1.23 (1.16, 1.30) | 0.77 (0.69, 0.84) |
| $\beta_W \times 10^{-6}$ | | 3.6 (2.85, 4.52) | 3.45 (3.22, 3.65) | 0.99 (0.76, 1.12) |
| $\beta_B \times 10^{-5}$ | | 1.71 (1.38, 2.22) | 1.08 (1.03, 1.13) | 0.41 (0.34, 0.52) |
| $p_C$ | | 0.38 (0.29, 0.47) | 0.34 (0.30, 0.38) | 0.41 (0.32, 0.48) |
| $p_D$ | | 0.26 (0.22, 0.31) | 0.21 (0.18, 0.24) | 0.03 (0.01, 0.08) |
| $\xi$ | | 3.17 (2.28, 3.92) | 3.63 (3.43, 3.92) | 2.78 (2.31, 3.19) |
| $\alpha_D$ | | | 0.71 (0.65, 0.74) | |
| $\epsilon_D$ | | | 0.61 (0.58, 0.63) | |
| $n_D$ | | | 3.25 (3.14, 3.34) | |
| $\alpha_R$ | | | | 13.5 (11.2, 16.7) |
| $\epsilon_R$ | | | | 3.45 (3.23, 3.71) |
| $n_R$ | | | | 5.92 (5.03, 6.86) |

To establish the ability of each model to capture the spatiotemporal characteristics of the observed outbreak, we now simulate our models using the posterior distribution for parameters summarised in Table 1. We use these simulations to determine the ability of each model at capturing both the temporal and the spatial epidemic profiles from the 2004 outbreak. This will provide an indication of whether the our modelling frameworks are able to mimic the fine-scale transmission dynamics between poultry farms in the country. The results are summarised in Fig 1.

**Figure 1:**
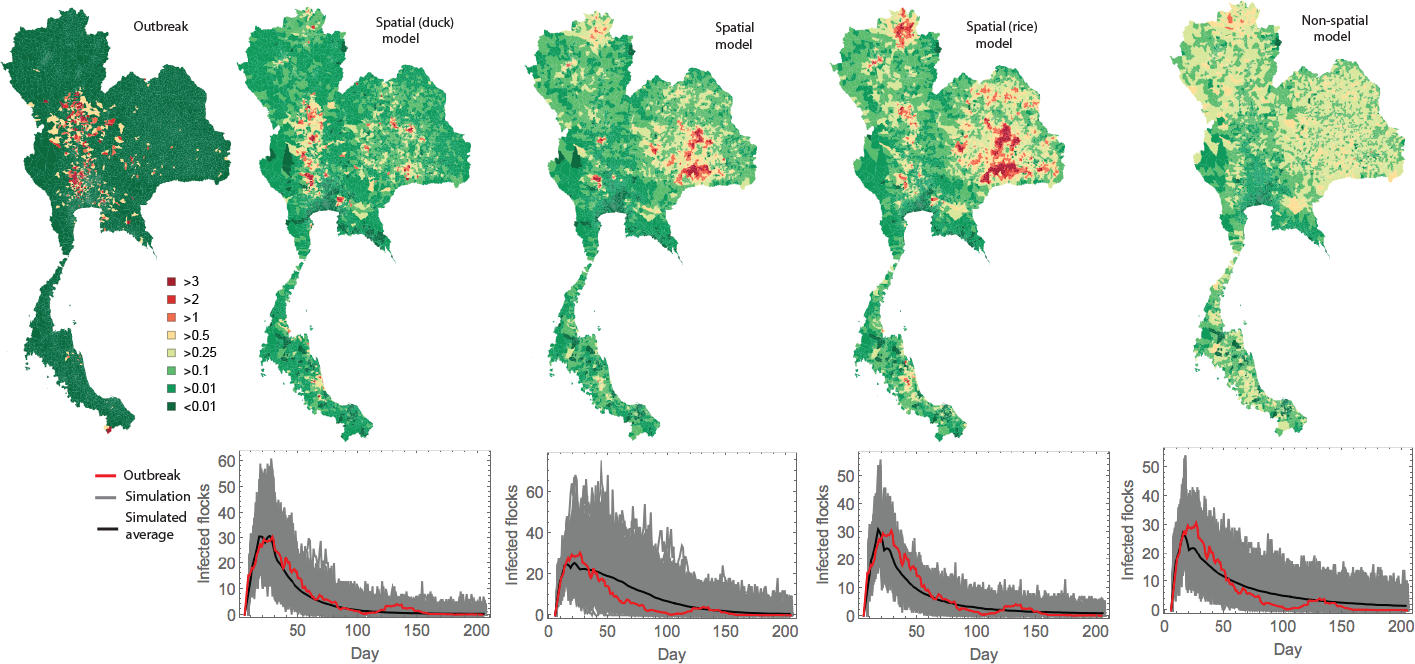
Predicted and observed outbreak distributions. Maps show the mean number of infected ocks in each subdistrict.

Model D appears to most accurately predict the spatiotemporal dynamics of the HPAI outbreak in Thialand (Fig 1, second column). For this model, spatial predictions of spread are much more representative of the true 2004 outbreak, with infected subdistricts predominantly in the centre and west of Thailand, in the region north of Bangkok. In addition, the temporal profile captures both the epidemic peak and the epidemic tail, providing supporting evidence of the significant role of the intensive duck farming industry in transmission of HPAI H5N1 in Thailand.

The spatial model, Model B, proved to be a poor predictor of geographical spread of the outbreak, with significant overestimates of infected farms in the east of the country and underestimates in the main epidemic hotspots. The model is also unable to accurately capture the temporal profile of the epidemic tail (Fig 1, third column). We see remarkably similar predictions for Model C (Fig 1, fourth column), though in this case overpredictions of spread in the east of Thailand are even greater, owing to the high densities of rice paddies in that part of the country. In addition, south eastern subdistricts of Thailand tend to have a higher average flock density, which will result in Models B and C overpredicting spread in these regions.

Random process model is unable to capture the spatial profile of the true outbreak, given that the model assumes random transmission characteristics. The model performs marginally better at predicting temporal behaviour, though slightly overestimates the epidemic peak and underestimates the number of farms infected in the epidemic tail (Fig 1, last column).

### 3.2. Sequential analysis

The model parameterisation that we have described in the section above determines parameter values by fitting the set of models to the entire observed spatiotemporal outbreak in Thailand. This gives us an understanding of the level of complexity required to capture HPAI epidemic dynamics. However, this approach only enables us to use our models retrospectively, to understand the model parameters and predictions of risk factors with the benefit of hindsight. However, it is also important to explore the predictive capacity of our models, so that we can assess whether the frameworks can be used to provide policy advice for ongoing epidemics.

In order to investigate the utility of our model predictions and control options in real time during outbreaks, we re-fit Model D only using outbreak data that would be available at particular points during the epidemic. The model is therefore parameterised using reported data up to day 10 and then sequentially on days 15, 20, 25, 30, 35, 50, 75, 100, 125, 150, 175 and 200 (the entire outbreak). We are interested in exploring how the prediction of the epidemiological parameters change through time. Our results are summarised in figure 2. As the observation time, *T*, increases, the estimated value of the background term *δ* decreases. Meanwhile, within subdistrict and between subdistrict transmission rates are both observed to initially increase as *T* increases, before decreasing again after the epidemic peak.

**Figure 2:**
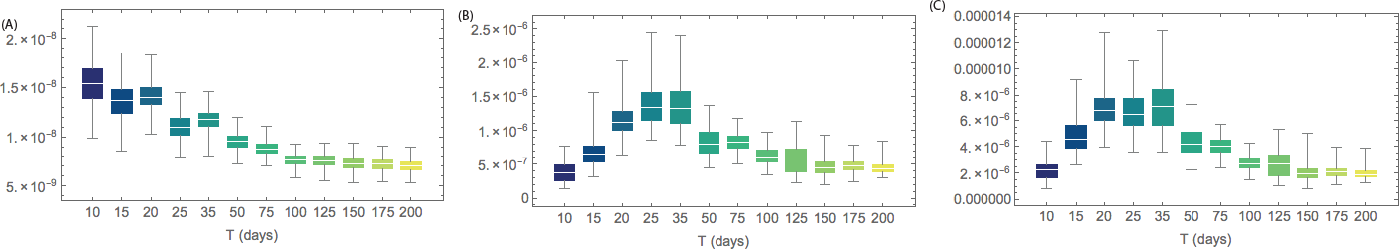
Distribution of fitted parameters for the spatial (ducks) model D: *δ* (A), *β*_*W*_ (B), and *β*_*B*_ (C). Sequential parameter estimation performed for outbreak data censored at time *T* days shown on *x*-axis.

When the model is only fitted to the first 10 days on the outbreak, we predict significant spread of the virus over large areas of the country (figure 3, second column). The model also dramatically overestimates outbreak size. This overestimate appears to be due to the parameterisation of the model on day 10 - the background term *δ* dominates (figure 2, fourth row) and the model predicts much lower within-subdistrict and cross border transmission (figure 2, fourth row). However, as the outbreak progresses, the model predicts that transmission is becoming more local, as seen by a reduction in the predictions of the value of *δ*. As the epidemic progresses, we see that the model performs much better at predicting the spatiotemporal profile of the outbreak. Indeed by day 30 (figure 3, third column), predictions of epidemic spread begin to be confined to regions where the true outbreak was observed. By day 50, the model is capable of producing accurate predictions of both the spatial and temporal profile (figure 3, fourth column), indicating that from this stage the ability of our modelling framework to generate robust predictions that could be used to inform policy in real time.

**Figure 3:**
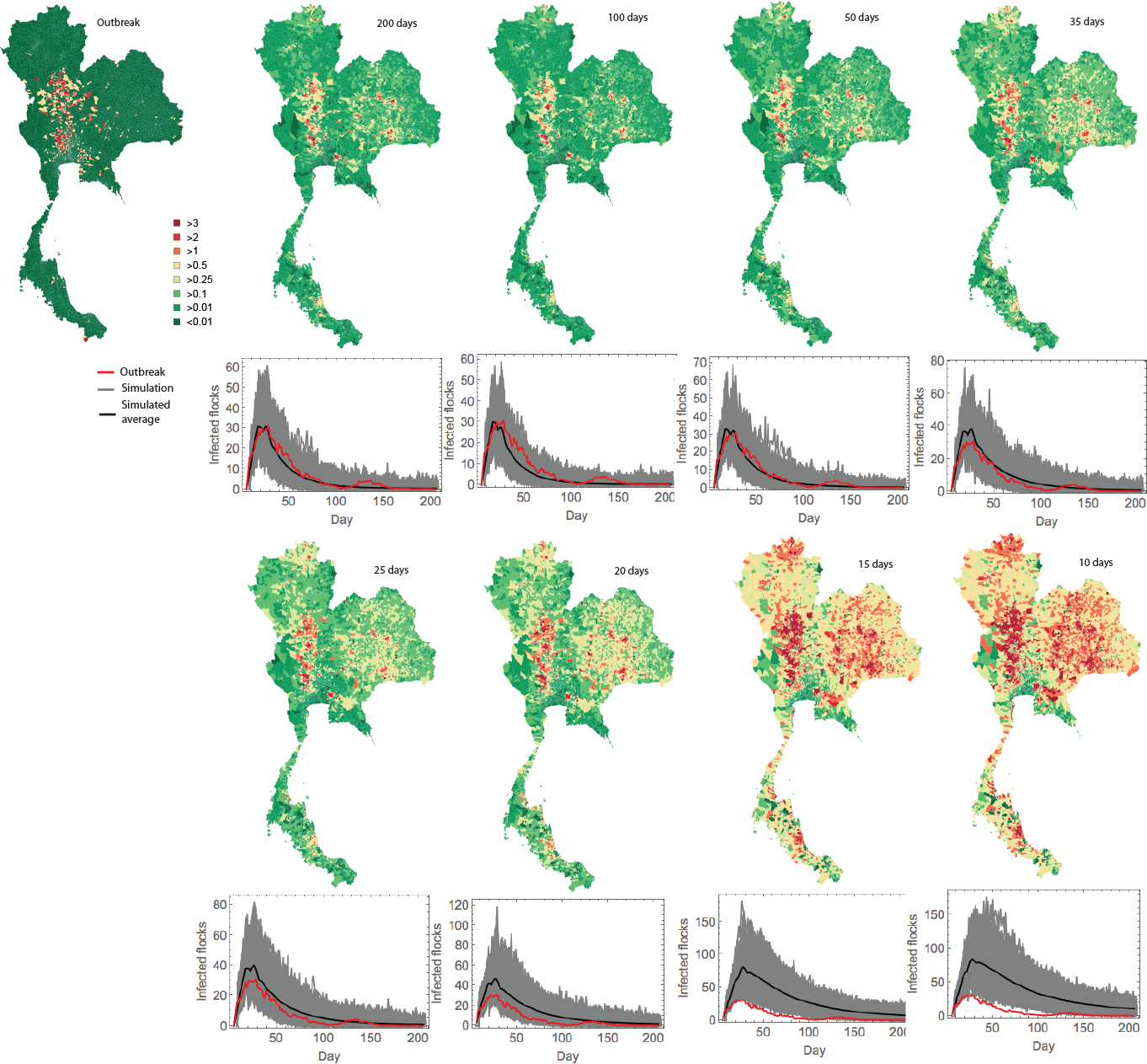
Predicted and observed spatial and temporal outbreak distributions. Each map show the mean number of infected flocks in each subdistrict as predicted by the model fitted at the given time in the outbreak. For each temporal profile, the observed 2004 outbreak is given by the red line, whilst the solid black line gives the mean model prediction.

### 3.3. Evaluating the effectiveness of local control

We now investigate the effectiveness of interventions at reducing the impact of H5N1 in Thailand. We use the four models and explore the impact of ring culling at radii from 0.25km to 5km around each infected premises and vaccination within infected subdistricts with capacities of 50%, 70% and 90% of all birds within the subdistrict. The results are summarised in (Fig 4*A*). We see a significant difference in predicted outbreak sizes and durations for the different control policies when we compare the predictions between the models. This suggests that, if we are interested in predicting the quantitative impact of a given intervention, it is crucial to select the most appropriate model. However, regardless of the differences in the predictions of epidemic size, all four models predict that large radius ring culling (5km) is the most effective strategy at minimising the total number of infected flocks (Fig 4*A*, top panel). This strategy results in a significant number of culled flocks and in fact if we are interested in minimising the total number of culled poultry flocks, high capacity vaccination proves to be the most effective policy (Fig 4*A*, middle panel). Models B, C and D predict that 5km ring culling results in the lowest outbreak duration across all the control strategies investigated. However, we see somewhat different results for our preferred spatial (duck) model - in this case there is no significant difference between a 5km ring culling policy and a vaccination strategy with a 90% capacity (Fig 4*A*, lower panel, Mann-Whitney test *p* = 0.14).

**Figure 4:**
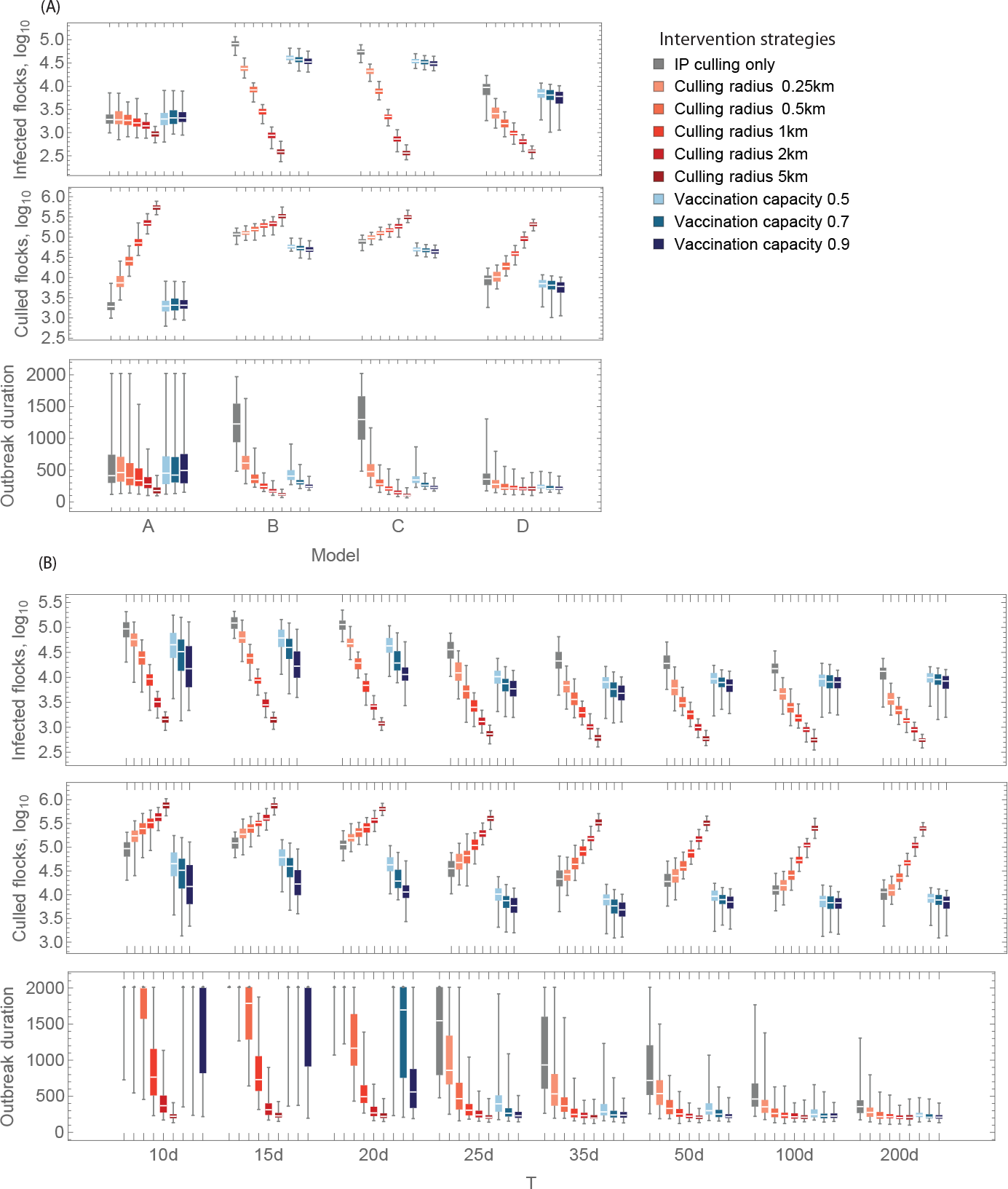
Effect of local controls. Projected number of infected flocks, number of culled flocks and duration of outbreak for (A) the four models, (B) for sequential parameter estimation using Model D with outbreak data censored at time *T* days.

Finally, we investigate the predictive ability of control policies for the preferred Model D. We therefore simulate our model from the start of the outbreak, using parameters that we have obtained on days 10, 15, 20, 25, 30, 35, 50, 100 and 200. We then explore the model predictions of the effectiveness of all competing ring culling and vaccination policies using model parameters obtained at each of these intervals. We observe that, even on day 10 when there is significant uncertainty in spatial spread parameters, Model D accurately predicts that 5km ring culling is the policy that minimises the total number of infected flocks (Fig 4*B*, top panel). Whilst the predicted epidemic sizes change for each control policy during the outbreak, the relative ordering of control policies remains consistent throughout. The same is true for total culled flocks, in that high capacity ring culling is always preferred (Fig 4*B*, middle panel).

However, if we are interested in minimising outbreak duration we see a somewhat different result. When the model uses only early outbreak data, predictions of epidemic duration are dramatically overestimated for IP only culling and all vaccination policies (Fig 4*B*, lower panel). The dynamics of the simulated outbreak is mostly dominated by the background *δ*, so even under a vaccination capacity of 90%, this gives a high enough infection pressure to sustain the outbreak for a long period of time. This overprediction is largely resolved by day 25 when vaccination is used, possibly owing to the more accurate predictions of the spatial extent of the disease at this point in the epidemic. We therefore conclude that it is important to determine the objective of control when deciding upon an intervention policy and that model predictions of the spread of the disease and the impact of interventions should be considered with caution during the early stages of any influenza epidemic.

## 4. Discussion

We have developed a set of mathematical models to investigate the level of complexity required to make accurate predictions regarding the spread of HPAI H5N1 in Thailand. When developing models, there is always a trade off between including enough model complexity to be able to capture the, often very complex, epidemiological and demographic characteristics that can lead to virus transmission, whilst at the same time keeping the model simple enough such that it is possible to parameterise. Here we explored that premise by considering four nested models of increasing complexity, where we accounted for random (non-distance based) spread, local spread of the disease, increased transmission owing to rice paddy density and increased transmission owing to presence of intensive duck farming.

Our results indicate that regions with a highly intensive duck farming industry are most likely to be infected during the H5N1 outbreak - our model that includes this factor provides the closest fit to the observational data from the 2004 outbreak. Our analysis reveals locations where the probability of finding infected animals is substantially higher, which can be used to identify specific areas where surveillance would be more beneficial than others in terms of increasing the probability of detecting the infection at early stages.

In addition, we have explored the ability of models of this nature to be used in real time during ongoing outbreaks. Many infectious disease models are used retrospectively to determine the risks associated with spread and the impact of control for previous outbreaks. Whilst this has value, it is also crucial to determine whether these models can be used in real time to advise policy makers during the course of an epidemic. Our results suggest that, during the early stages of HPAI outbreaks, there is significant uncertainty, owing to partial reporting of cases and potential for undetected infections over large areas. Models such as the one we describe here therefore only have limited predictive power in the very early stages of HPAI epidemics. However, as more data are accrued, the uncertainty in predictions decreases, such that after the first few weeks, we are able to accurately predict both the size and the spatial extent of the outbreak. This is highly beneficial in terms of being able to inform targeted surveillance policies to improve detection.

Despite this uncertainty, the models perform significantly better in determining the optimal control policies that should be deployed, even in the very early stages of the outbreak. This is promising, in that, despite over predictions of spread at epidemic onset, models are able to inform how to target interventions to reduce the risk of spread in the future. The caveat to this is the uncertainty in the impact of some control policies in the early stages of outbreaks - whilst the model accurately predicts that high radius ring culling is the preferred policy even when only using the first 10 days of the outbreak to parameterise the model, the duration of outbreaks is significantly overpredicted for IP only culling and vaccination policies.

For the future, it is important to explore mechanisms to improve the predictive power of infectious disease models during the early stages of disease outbreaks. It is simply not possible for policy makers to wait for uncertainty to resolve after the first few weeks before employing an intervention and therefore seeking methods to reduce uncertainty in model predictions at epidemic onset is vital. One area for future research would be to use data from previous outbreaks to provide more informed priors for epidemiological parameters. This would enable us to explore whether this can better establish the likelihood of spread in the first few days of a new epidemic when the outputs of models could potentially have the most significant impact upon reducing the spread of the disease.

## 5. Supplementary Material

### 5.1. Data

The demographic data describing poultry industry in Thailand during the 2004 outbreak has the following characteristics: number of subdistricts: 7,416; number of flocks: 3,303,160; number of chicken: 180,725,929; number of ducks: 19,745,049. This gives that the national poultry density during the outbreak was 395 *birds/km*^2^.

### 5.2. Mathematical models

We formulate four mathematical models with increasing complexity: a random process model (Model A), a spatial model (Model B), a spatial model incorporating rice (Model C), and a spatial model incorporating duck intensity (Model D).

Any district *k* is described by a number of flocks, *F*_*k*_, number of susceptible flocks at time *t*, *S*_*k*_(*t*), and number of in infectious flocks at time *t*, *I*_*k*_(*t*). Then any flock *i* in a subdistrict *k* is described by a number of chicken, *NC*_*k, i*_ and number of ducks, *ND*_*k,i*_. For further calculations, we normalise number of chicken and ducks as follows:

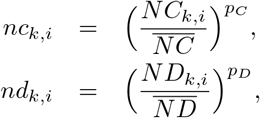

where power laws *p*_*C*_ and *p*_*D*_ account for virus transmission differences in the two species.

We can write a general expression for the force of infection to a susceptible flock, *k*:

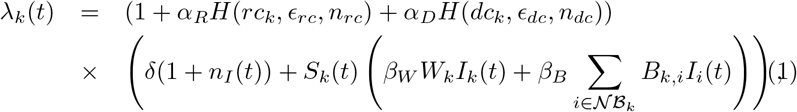

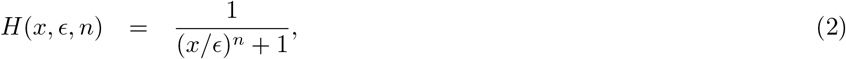

Here *rc*_*k*_ is the fraction of the area of the subdistrict occupied by rice paddies; the intensity of duck industry is formalised as *dc*_*k*_ = log_10_ (*max*_*i*=1‥*F*_*k*__ (*N D*_*k,i*_) + 1); *n*_*I*_ (*t*) is the number of infected flocks at time *t*, whilst *β*_*W*_ and *β*_*B*_ are the rates of spatial transmission within and between subdistricts, whilst *W*_*k*_ and *B*_*k,i*_ are the rates of within-subdistrict transmission and local cross-border transmission. The term *δ* is spatially independent and allows transmission between any pair 410 of subdistricts. We refer to *δ* from here onwards as the “background” term.

The rate of within-subdistrict transmission for subdistricts with a number of flocks *F*_*k*_ > 1 is given by

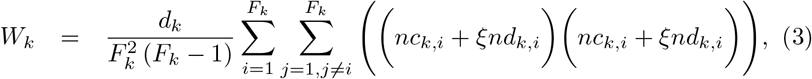

where *d*_*k*_ is the density of flocks (i.e. number of flocks divided per subdistrict area) and *F*_*k*_ is a number of flocks in a subdistrict *k*.

The rate of local cross-border transmission between subdistricts with a number of flocks *F*_*k*_ > 0 and neighbouring subdistricts 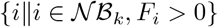 is given by

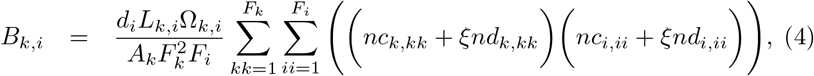

where *L*_*k,i*_ is the length of a shared boundary between neighbour districts *k* and *i* and *A*_*k*_ is the area of subdistrict *k*.

The term Ω_*k,i*_ is a scaling factor for between-subdistrict transmission informed by the mean distance between neighbouring subdistricts taking into account the size of each subdistrict, following the approach of Buhnerkempe et al. 2014 such that:

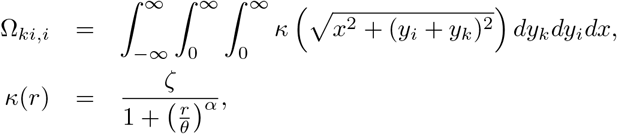

where *α* and *θ* are the shape and scale parameters of the distance-dependent local spread kernel respectively (Buhnerkempe et al. 2014). The normalisation constant *ζ* is defined such that

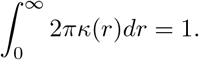

### 5.3. Bayesian approach for parameter estimation

We use a Bayesian approach to fit the meta-population model to the outbreak data (Jewell et al. 2009). The likelihood function has the following expression:

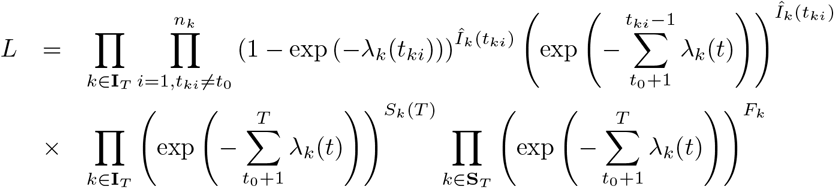

where *n*_*k*_ is the total number of infection events in a subdistrict *k* and *t*_*ki*_ is the time of the *i*^*th*^ infection event in a subdistrict *k*. We introduce 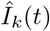 to be the number of flocks that have been infected at time *t*, and note that this is different from *I*_*k*_ (*t*) (i.e. the number of infectious flocks at time *t*). For each of the parameters, we update the parameter value using a random walk Metropolis algorithm with a Gaussian proposal: *θ*\* ~ *N*(*θ*, *σ*), where *θ* and *θ*\* are parameter vectors, and *σ* is a covariance matrix.

The proposed set of updated parameters *θ*\* is then accepted with probability

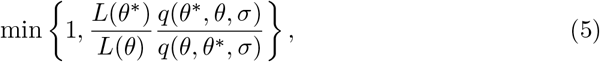

where *q*(*x*, *y*, *s*) is a normal PDF value at *x* with mean *y* and covariance matrix *s*.

As a pre-emptive culling policy within a 1km radius of flocks reporting infection was introduced during the outbreak, we simulate this by reducing the number of susceptible flocks as a fraction given by the ratio of the area of a circle with 1km radius to the total area of the subdistrict in question. We assume that there was no re-population of any flocks after culling prior to the end of the epidemic.

### 5.4. Model simulation

Posterior samples of parameters are used to generate predicted outbreaks. At every time *t* of the simulation (time step Δ*t* = 1 day), every susceptible flock *k* has a probability of becoming infected:

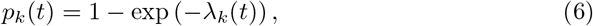

where the infection pressure depends on a particular model. We start the outbreak by seeding infection in the six subdistricts in Thailand that had reported infection during the first three days of the outbreak. The seed cases were the same for all simulations. The outbreak was assumed to end after seven days without new infected cases (or alternatively the simulation was terminated after 2000 days).

#### Modeling Intervention strategies

With vaccine capacity *γ*, the demographic factors on the infection pressure then are reduced so that:

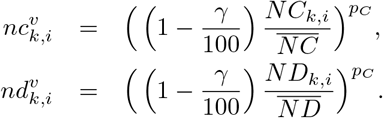

#### Financial disclosure

MJT and RR are supported by the Engineering and Physical Sciences Research Council and the Biotechnology and Biological Sciences Research Council [grant numbers EP/P511079/1 and BB/K010972/4]. XX, MG and MJT are supported by the National Institutes of Health (NIH grant 1R01AI101028-02A1). The funders had no role in study design, data collection and analysis, decision to publish, or preparation of the manuscript.

### Data availability

Data are available from the Department of Livestock Development in Thailand.

## Acknowledgments

We thank Will Probert and Ed Hill for useful comments on this manuscript.

## Author Contributions

The author(s) have made the following declarations about their contributions: Conceived and designed the experiments: MT, RR, CJ. Performed the experiments: RR, MT. Analyzed the data: RR, WT. Contributed reagents/materials/analysis tools: TB, MG, XX, GZ, WT. Wrote the paper: RR, MT, MK, TB, MG, GZ, XX, CJ, WT.

